# On the diversity of F_420_-dependent oxidoreductases: a sequence- and structure-based classification

**DOI:** 10.1101/2020.08.24.261826

**Authors:** María Laura Mascotti, Maximiliano Juri Ayub, Marco W. Fraaije

## Abstract

The F_420_ deazaflavin cofactor is an intriguing molecule as it structurally resembles the canonical flavin cofactor, although biochemically behaves as a nicotinamide cofactor. Since its discovery, numerous enzymes relying on it have been described. The known deazaflavoproteins are taxonomically restricted to Archaea and Bacteria. The biochemistry of the deazaflavoenzymes is diverse and they exhibit some degree of structural variability as well. In this study a thorough sequence and structural homology evolutionary analysis was performed in order to generate an overarching classification of all known F_420_-dependent oxidoreductases. Five different superfamilies are described: Superfamily I, TIM-barrel F_420_-dependent enzymes; Superfamily II, Rossmann fold F_420_-dependent enzymes; Superfamily III, β-roll F_420_-dependent enzymes; Superfamily IV, SH3 barrel F_420_-dependent enzymes and Superfamily V, 3 layer ββα sandwich F_420_-dependent enzymes. This classification aims to be the framework for the identification, the description and the understanding the biochemistry of novel deazaflavoenzymes.

## Introduction

F_420_ is a naturally occurring deazaflavin cofactor in which the N5 atom of the isoalloxazine ring is substituted by a C atom and has an 8-hydroxyl moiety, compared to the canonical flavin cofactors FMN and FAD. It was first isolated almost 50 years ago by Cheeseman *et al* [1] and its structure solved shortly after [2]. F_420_ is an obligate two-electron hydride carrier and it shows a low standard redox potential (−340 mV), which resembles that of nicotinamide cofactors (−320 mV) rather than that of flavins (−220/−190 mV) [3]. The first reports on F_420_-using enzymes were related to methanogenesis in archaeal species [4, 5]. For a long time these were considered as unusual proteins. Recent research has demonstrated that they are actually widespread across Archaea and Bacteria [3]. The species from the euryarchaeota phyla, *Methanosarcina* spp, *Methanothermobacter* spp and *Archaeoglobus fulgidus* are among the most frequently investigated in Archaea. In Bacteria, research has been focused in the actinobacteria genus *Mycobacterium*. Notably, in *M. tuberculosis* the F_420_-dependent enzymes have been associated to its pathogenicity [6]. The restricted domain distribution of the F_420_ cofactor and its connection to anaerobic metabolism, highlight its status as a relic from the origin-of-life world.

A number of F_420_-dependent enzymes have been classified according to their three-dimensional fold into three groups: the luciferase-like monooxygenases (LLM), the pyridoxamine-5'-phosphate oxidases (PNPOx), and the deazafavin-dependent nitroreductases (DDN) [7]. Although this classification was based in structural homology, it should be noted that the PNPOx and DDN representatives display the same split barrel fold. While all known DDNs rely exclusively on F_420_, LLM and PNPOx members show dependence on other flavin cofactors as well [8, 9]. The aflatoxin degrading F_420_-dependent reductases from actinomycetales were discovered and characterized recently [10]. These were described as F_420_-dependent reductases (FDR-A and FDR-B) and are homologous to members of the PNPOx family [11]. Later, it was proposed that FDRs should be instead referred to as flavin/deazaflavin oxidoreductases (FDORs A and B). The FDOR-A group includes exclusively F_420_-depedent enzymes, while the FDOR-B encompasses deazaflavoenzymes as well as enzymes using FMN, FAD, and heme cofactors [12]. Except for those enzymes utilizing F_420_H_2_ in the reduction of metabolites, there are many known proteins that use the deazaflavin cofactor for other reactions. Among them are oxidoreductases that can shuttle a hydride between nicotinamide and F_420_ (FNOs) [13, 14], oxidases that use F_420_ coupled to FMN to reduce dioxygen (FprA) [15], and dehydrogenases employing F_420_ associated to methylene-H_4_MPT cofactor [16]. Additionally, other redox enzymes in anaerobic metabolism such as the [NiFe]-hydrogenases [17] and the thioredoxin reductases [18] depend on the F_420_ cofactor and show unique structural features. Therefore, considering the increasing number of characterized F_420_-dependent enzymes displaying not only different biochemistries but also a variety of structural topologies, a structure-based classification of deazaflavoproteins would be valuable.

Enzyme classification can be performed either on the basis of functionality a ‒*e.g.*: the chemical reaction they catalyze [19, 20]-, or on the basis of the evolution [21]. One of the main drawbacks of the first strategy is that it relies on features lacking a common evolutionary origin. Although these phenetic classifications are a very useful way of organizing protein knowledge, problems arise when investigating the underlying determinants of enzymes’ functionality. Classifications based on overall similarities also perform poorly predicting functionalities of new enzymes and when the number of items to classify increases constantly over time [19]. Molecular evolution allows understanding how modern enzymes work as they do [22]. Therefore, the classifications based on it constitute a framework to infer the activity of new enzymes and to explore its physico-chemical and biological determinants.

In this work all enzymes that use the deazaflavin cofactor F_420_ were comprehensively analyzed from the structural homology perspective. Five different superfamilies embracing the whole enzymatic diversity of F_420_-dependent oxidoreductases recognized at the moment have been identified. Besides, the evolutionary history of these superfamilies is reported and the trends in the cofactor utilization are analyzed.

## Results and Discussion

A molecular evolution analysis was conducted aiming to integrate the current biochemical knowledge on F_420_-dependent enzymes. All currently known enzymes that use F_420_/F_420_H_2_ as cofactor in oxidation/reduction processes were included. A shared structural fold was considered as the first sign of a common evolutionary origin [21]. Regarding the nomenclature, the names of the different groups were conserved whenever possible in order to ease the use of the classification presented here.

Initially, the strategy consisted in mining structural databases to get all enzymes using F_420_ as a cofactor. The 50 collected structures (out of 171718 PDB entries scanned) belong to 24 different oxidoreductases. By analyzing the domain topology and architecture, these enzymes were assigned to four different evolutionary-independent units. In that way, four different unrelated folds were identified among the gathered structures. For those enzymes with no structure available [18], a second dataset was built and the domain topology predicted on the basis of CATH [23]. From this analysis, one extra distinct fold was identified. Structure-based alignments were constructed when possible for each of the identified groups and the evolutionary relationships were inferred. Next, homology searches and HMM profiling were employed to find close and distant homologs and sequence-based phylogenies were constructed (see Methods section). All the evolutionary and structural information obtained was integrated with the biochemical data. This step-wise analysis revealed the existence of five well-defined deazaflavoenzyme superfamilies (I-V) (Table 1 & Table S1). These are presented and described in the following sections.

**Table 1.**
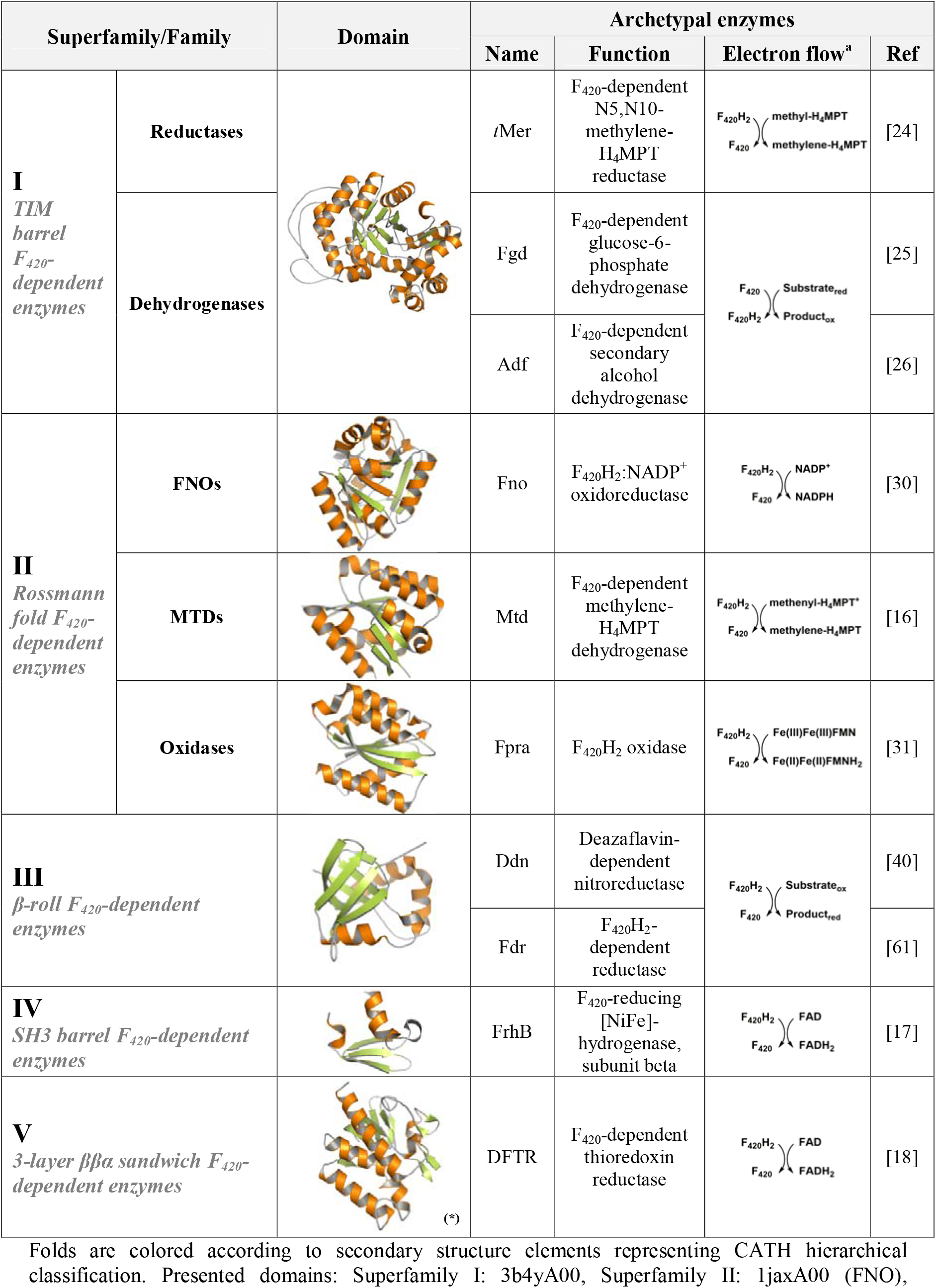
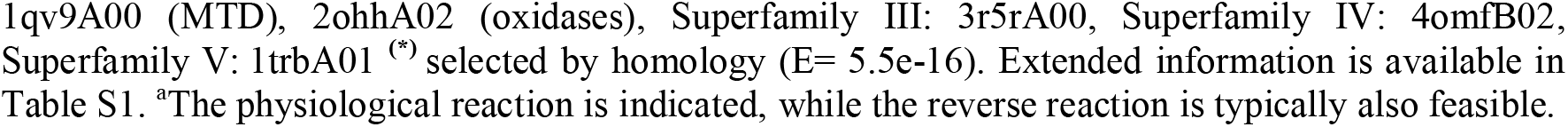
The deazaflavoenzyme superfamilies.

| Superfamily/Family |  | Domain | Archetypal enzymes |  |  |  |
| --- | --- | --- | --- | --- | --- | --- |
|  |  |  | Name | Function | Electron flow<sup>a</sup> | Ref |
| **I**<br>*TIM barrel*<br>*F<sub>420</sub>-dependent enzymes* | Reductases |  | *t*Mer | F<sub>420</sub>-dependent N5,N10-methylene-H<sub>4</sub>MPT reductase |  | [24] |
|  | Dehydrogenases |  | Fgd | F<sub>420</sub>-dependent glucose-6-phosphate dehydrogenase |  | [25] |
|  |  |  | Adf | F<sub>420</sub>-dependent secondary alcohol dehydrogenase |  | [26] |
| **II**<br>*Rossmann fold F<sub>420</sub>-dependent enzymes* | FNOs |  | Fno | F<sub>420</sub>H<sub>2</sub>:NADP<sup>+</sup> oxidoreductase |  | [30] |
|  | MTDs |  | Mtd | F<sub>420</sub>-dependent methylene-H<sub>4</sub>MPT dehydrogenase |  | [16] |
|  | Oxidases |  | Fpra | F<sub>420</sub>H<sub>2</sub> oxidase |  | [31] |
| **III**<br>*β-roll F<sub>420</sub>-dependent enzymes* |  |  | Ddn | Deazaflavin-dependent nitroreductase |  | [40] |
|  |  |  | Fdr | F<sub>420</sub>H<sub>2</sub>-dependent reductase |  | [61] |
| **IV**<br>*SH3 barrel F<sub>420</sub>-dependent enzymes* |  |  | FrhB | F<sub>420</sub>-reducing [NiFe]-hydrogenase, subunit beta |  | [17] |
| **V**<br>*3-layer ββα sandwich F<sub>420</sub>-dependent enzymes* |  | (*) | DFTR | F<sub>420</sub>-dependent thioredoxin reductase |  | [18] |

Folds are colored according to secondary structure elements representing CATH hierarchical classification. Presented domains: Superfamily I: 3b4yA00, Superfamily II: 1jaxA00 (FNO),

### Superfamily I: TIM barrel F_420_-dependent enzymes

This group is described by the alpha-beta barrel domain and includes the so-called LLMs. Two kinds of F_420_-dependent oxidoreductases are found here: the methylene-H_4_MPT reductases (MERs) [24] and the dehydrogenases, represented by the F_420_-dependent glucose-6-phosphate dehydrogenases (FGDs) (Table 1) [25]. Under physiological conditions the MERs consume F_420_H_2_ and the FGDs generate it. Regardless of their apparent opposite biochemistry the reversibility of their activities has been demonstrated [26].

The sequence-based phylogeny strongly indicates a single common origin for all deazaflavoenzymes in this superfamily (TBE= 0.96) branching out from FMN-dependent enzymes (Fig. 1 & Fig. S1). Two well-defined clades of deazaflavoenzymes are observed. One includes the MERs, and we have named this the *reductase* clade (TBE= 0.95). The other is populated by the FGDs and alcohol dehydrogenases, thus referred as the *dehydrogenase* clade (TBE= 0.99). The FMN-dependent sister clade includes proteins such as the bacterial alkanesufonate monooxygenase (ssuD, PDB: 1m41) [27] and the nitrilotriacetate monooxygenase (NTA_MO, PDB: 3sdo), as well as the well-known bacterial FMN-dependent luciferases (*e.g.*: LuxB, PDB: 1luc) [28]. The topology suggests that cofactor specificity (FMN or F_420_(H_2_)) has played a major role as selective pressure in the evolutionary history of this superfamily. Recently we proposed that the cofactor switching event from FMN to F_420_ occurred *via* a gene duplication event resulting in F_420_ usage [29]. The taxonomic distribution shows a mixed arrangement of bacterial and archaeal species with predominance of facultative anaerobes and methanogens.

**Figure 1.**
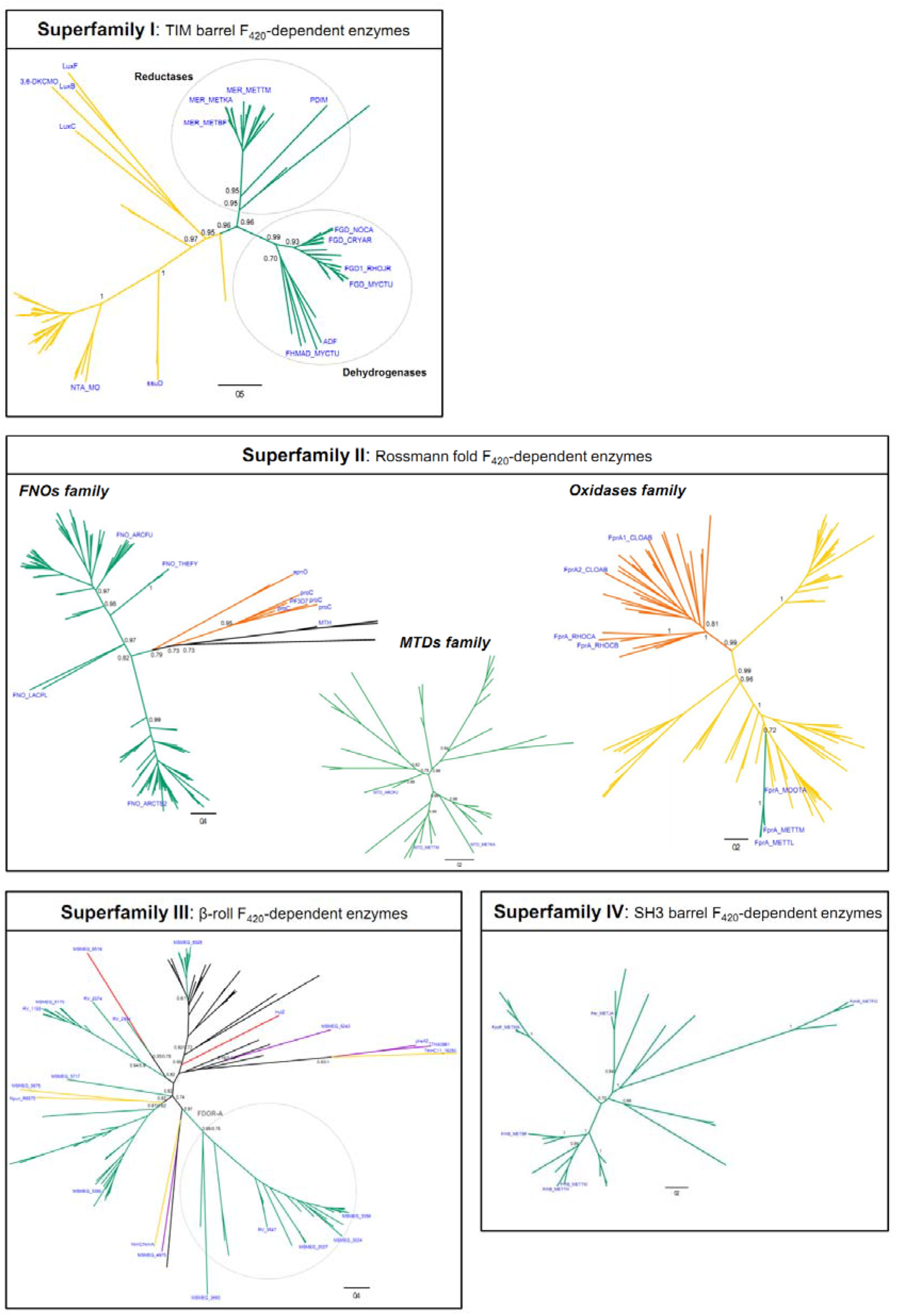
Phylogenetic trees of superfamilies I-IV. Sequence-based phylogenies are presented unrooted. Support values (TBE) corresponding to major divergences are indicated at the nodes. For superfamily III also PP values from Bayesian inference are given next to the TBEs. Names (in blue) are given for those enzymes that have been experimentally characterized. Cofactor specificities are indicated with the color of the branches as follows: F_420_ (green), FMN (yellow), NAD(P)H (orange), FAD (purple) and heme (red). Unknown cofactor specificity is represented with black branches. For fully annotated trees please see the Supporting information (Figures S1 to S4).

### Superfamily II: Rossmann fold F_420_-dependent enzymes

Superfamily II includes deazaflavoenzymes sharing the classic 3-layer αβα sandwich topology, commonly known as Rossmann fold. Members with known biochemical features vary considerably in structure and function, and comprise the F_420_H_2_-NADP^+^ oxidoreductases (FNOs) [30], the F_420_-dependent methylene-H_4_MPT dehydrogenases (MTDs) [16], and the F_420_H_2_ oxidases (FprAs) [31]. The latter show a unique multidomain architecture, since they are N-terminally fused to a □180-amino acid β-lactamase domain (PF00753, CATH 3.60.15.10). Despite the evidenced structural differences, catalysis is similar as the hydride transfer proceeds from F_420_H_2_ to a second molecule/cofactor such as a nicotinamide (FNOs), tetrahydromethanopterin (MTDs) or a flavin (FprAs) [31] (Table 1).

In spite of some structural variations of the central common fold, a structure-based phylogeny could be constructed (Fig. S2A). MTDs and oxidases emerge as clear monophyletic groups (BS=97 and BS=100, respectively). FNOs, though sharing ancestry, form a poorly supported paraphyletic group. Considering the topology of the structural phylogeny plus the sequence similarity among each group, we propose the existence of three families inside the superfamily. These are coined FNOs, MTDs, and oxidases (Table 1). Phylogenies were constructed for each of them, revealing some shared and other unique evolutionary paths (Fig. 1 & Fig. S2).

The FNOs family includes all known F_420_H_2_-NADP^+^ oxidoreductases. Phylogenetic analysis shows that all members of this family derive from a single ancestor (TBE= 0.82) and are distributed in two groups (Fig. 1 & Fig. S2B). This clear splitting does not respond to any taxonomic or structural features and thus should be further investigated. The FNO-type sequences are 200-220 amino acids in length, show single domain architecture (PF03807, CATH 3.40.50.720) and use F_420_ coupled to NADP^+^. FNOs are homologous to some NAD(P)H-dependent reductases which encode for an extra C-terminal domain of 40-50 amino acids length. These reductases are early diverging in the phylogeny and do not use a deazaflavin cofactor coupled to the nicotinamide (Fig. 1). Among them, the pyrroline-5-carboxylate reductases (forming a monophyletic group of 5 sequences, TBE= 0.95) [32] and the D-apionate oxidoisomerase (apnO, PDB: 5t57) [33] are found (Fig. S2B). The topology of the tree strongly suggests that the use of nicotinamide cofactor is the ancestral feature whereas the utilization of F_420_ arose later in evolution. Taxonomic distribution of FNOs seems to be enriched in anaerobic or facultative anaerobic species either from Bacteria or Archaea.

The MTDs family includes the known F_420_-dependent methylenetetrahydromethanopterin dehydrogenases (Fig. 1, Fig. S2C). This family is composed solely of F_420_-using enzymes that display an exclusive structural domain (PF01993, CATH 30.40.50.10830). The taxonomic distribution of MTDs is strongly biased towards methanogenic archaea belonging to the euryarchaeota phylum [16, 34]. Likewise, these enzymes are present in archaeoglobi species [35] which, despite being sulfur-metabolizing organisms, encode a nearly complete set of genes for methanogenesis [36]. A few eubacterial-derived proteins are also observed. However, none of these sequences have been experimentally characterized.

The family including the F_420_H_2_ oxidases (FprA) shows another different history (Fig. 1 & Fig. S2D). For a more inclusive denomination, we called this family as the oxidases. The two-domain architecture (lactamase B [pfam 00753]- flavodoxin [pfam 00253]) defining this group is shared by various other redox enzymes such as nitric oxide reductases [37] and flavodiiron proteins [38]. The evident difference among these FMN-dependent enzymes is the electron donor preference, which can be F_420_H_2_ or NADH. For those working as nitric oxide reductases no other cofactor than FMN is required. Phylogenetic analysis shows that F_420_ usage is restricted to one particular clade (TBE= 1) containing all known FprAs, embedded into a large group of FMN-dependent enzymes (Fig. 1 & Fig. S2D). On the other hand, those enzymes reportedly relying on NADH as electron donor form a monophyletic group (TBE= 1) which has a sister clade of putative FMN-dependent homologs. Therefore, the ancestral feature seems to be the usage of FMN, while two later independent events yielding F_420_H_2_ and NADH specificities have taken place. Remarkably, the utilization of F_420_ in this family seems to be restricted to archaeal species [15, 31], while NADH is used by bacteria [38, 39]. However, this statement should be considered cautiously as very few members of the family have been experimentally characterized.

### Superfamily III: β-roll F_420_-dependent enzymes

This superfamily encompasses F_420_-dependent reductases of great biochemical diversity: the well-known DDNs [40] and the F_420_H_2_-dependent reductases [11], also referred as FDOR-A & B [12]. All enzymes display an arrangement of antiparallel beta sheets commonly called split barrel fold, which actually falls within the mainly β-roll architecture according to CATH [23]. The well-known FMN-dependent PNPOx [9] and other enzymes displaying specificity for FAD or heme cofactors are related to the sequences in this group. The available protein structures allowed us to construct a structure-based phylogeny with the purpose of unveiling if clustering corresponds to cofactor specificity. The topology of the obtained tree revealed that the superfamily has no structurally imposed cofactor preference, as an interleaved distribution is observed (Fig. S3A). This was further confirmed by a sequence-based phylogenetic analysis which resulted in the same clade distribution. By constructing a robust dataset ‒*via* ensuring diverse taxonomic representation and lowering redundancy- the scattered cofactor dependence became clear (Fig 1 & Fig. S3B). The tree shows some hard polytomies even after inferring the phylogeny by Maximum Likelihood and Bayesian methods. However, well-supported clades could be identified and trends in cofactor usage became clear. These are discussed as follows.

All sequences previously characterized as FDOR-A [12] (MSMEG_3660, 2027, 3004, 3356 and Rv3547) sharing the use of F_420_ molecule as cofactor, form a monophyletic group (TBE/PP= 0.95/0.75). However, the cofactor preference for F_420_, FMN, FAD and heme is interspersed in the rest of the tree. It becomes evident that the FDOR-B (B) members [12] do not form a monophyletic group. The clade comprising the aflatoxin degrading enzyme MSMEG_3380 (B) includes the early diverging FMN-dependent enzymes Npun_R6570 from *Nostoc punctiforme* and MSMEG_5675 with very good confidence (TBE= 0.82). Similarly, a distinct clade is observed containing the F_420_-dependent enzymes Rv1155 (B), MSMEG_5170 (B), Rv2074 (B), Rv2991 and the heme-dependent enzyme MSMEG_6519 (TBE=0.82). This irregular distribution becomes even more evident in the clade including the F_420_-dependent MSMEG_6526 and the heme storage protein HutZ (TBE= 0.89). Among this group an early diverging set of FAD- and FMN-dependent reductases is found. These are the MSMEG_5243, the reductase component of the aromatic hydroxylases pheA2 and TTHA0961 [41, 42], and the small reductase component of a styrene monooxygenase TthHC11_19250 [43]. From this analysis, it is evident that cofactor preference is not the trait selected during evolution. This also echoes in the flexibility of the cofactor binding site, which allows certain degree of promiscuity as some reductases seem to be able to utilize both F_420_ and FMN [44]. This interspersed distribution imposes implications for applied enzymology, as cofactor specificity of new enzymes can be hardly predicted from sequence and/or structure for this superfamily. It has to be experimentally determined as it has been previously shown [12]. Finally, it should be noted that although most of the research has been devoted to *Mycobacterium* species, proteins from other bacteria as well as Archaea are observed among all clades (Fig. S3B).

### Superfamily IV: SH3 barrel F_420_-dependent enzymes

Superfamily IV is formed exclusively by deazaflavoenzymes. These are the F_420_-reducing [NiFe]-hydrogenases, a complex of three subunits (FhrABG) involved in the oxidation/reduction of F_420_ in the methanogenesis pathway in Archaea [17, 45, 46]. Subunit B (FhrB) contains a FAD molecule, a [4Fe-4S] center and the F_420_ binding site. It displays a characteristic SH3 barrel topology at the N-terminal region. This subunit is devoted to the reduction of F_420_ at the expenses of electrons transferred from a flavin/ferredoxin system (Table 1). Despite the lack of many 3D structures, a few homologs have been characterized in detail. When analyzing their predicted topology they all display the same structural domain. Fpo-F [47], Fhr-B [46], F_420_ sulfite dehydrogenase (FSR) [48] and Hdr (heterodisulfide reductase) [49] populate this superfamily. All sequences in this superfamily belong to the Archaea domain exclusively (Fig. 1, Fig. S4).

### Superfamily V: 3-layer ββα sandwich F_420_-dependent enzymes

Superfamily V has only a single biochemically characterized member with no solved 3D structure, namely the deazaflavin-dependent flavin-containing thioredoxin reductase (DFTR) from *Methanocaldococcus jannaschii* (Mj-TrxR) [18]. This enzyme has a predicted fold similar to the NAD(P)H-dependent thioredoxin reductases (NTR) consisting in a three-layer sandwich. However, the classic nicotinamide cofactor is replaced by F_420_H_2_ as the electron donor to reduce the bound FAD (Table 1). The sequence analysis does not reveal any unique feature accounting for such a change in the cofactor specificity, rather than a few changes in the predicted NAD(P)H binding regions. From the evolutionary history [18], it seems clear that DFTR belongs to the disulfide oxidoreductase family of flavoenzymes. However, as the majority of the close homologs have not been experimentally characterized it is hard, if not impossible, to make assumptions on the electron donor specific distribution across the superfamily. It has been proposed that DFTRs would be only present in the methanococci Archaea class replacing the NTRs [50]. If that were the case, the utilization of F_420_H_2_ would be a derived feature of the protein family. However, a deeper understanding of this superfamily is strongly contingent on the discovery of new deazaflavoenzymes displaying this fold.

## Conclusion

A global classification of the oxidoreductases that employ the deazaflavin F_420_ cofactor is proposed on the basis of the evolutionary analysis of available sequences and structures. This clustering may allow identifying novel F_420_-dependent enzymes as well as to disclose the origin and extent of the functionalities associated with F_420_ cofactor. Furthermore, this study provides clues on how F_420_-dependent enzymes are evolutionary related to other redox enzymes that rely on cofactors like heme, flavin, and nicotinamide.

## Methods

### Evolutionary clustering of F_420_-dependent enzymes

PDB files from all F_420_-dependent enzymes were collected from PDBsum [51]. Domain architecture was investigated using the hierarchical criteria from CATH [23] and/or pfam [52]. The InterPro database was also investigated [53] (last accession to all databases: 12 august 2020). Structures were specifically clustered according to their domain topology and architecture as the definitive criteria for a shared evolutionary origin. Structural alignment of each group was performed in PROMALS3D [54].

### F_420_ protein superfamilies profiling

For each group (superfamily), sequence datasets were constructed by homology searches using as queries the sequences of the enzymes previously identified (*vide supra*). Blastp was conducted using non-redundant protein sequences database and specifying the taxonomy (Archaea or Bacteria). The 250/500 first hits were collected on each search (E-value ≥ 1e-9). HMM profiling was also conducted in HMMER using UniprotKB database and the first 250/500 hits were collected. Enzymes sharing the same structural domain but not using the F_420_ cofactor were also used as queries in the homology searches. Retrieved sequences were included in the datasets to ensure clustering unbiased by the data collection. For superfamily II pfam available datasets were collected in complete form as well. Raw datasets contained >1000 seqs for each superfamily. MAFFT v7 was employed to build multiple sequence alignments (MSAs) [55]. Redundancy was removed (cut-off = 80% identity) with CD-HIT [56]. MSAs were visually inspected and single sequence insertions/extensions trimmed. Substitution models and alignment parameters were calculated in ProtTest 3.4.2 [57]. Phylogenies were constructed by the maximum likelihood inference method implemented in PhyML 3.1 [58] or RaxML 8.2.12 [59] with 100/500 bootstraps (BS), respectively. Transfer bootstrap values (TBE) were obtained in BOOSTER [60]. For superfamily III, Bayesian inference was also conducted in Mr. Bayes 3.2.6 running 2000000 generations until convergence < 0.2 was reached. FigTree 1.4.2 was employed to visualize and edit the trees.

## Author Contributions

MLM & MWF conceived the original idea. MLM designed the research and performed all the analyses. MLM, MJA & MWF interpreted the data. MLM proposed the classification system and wrote the paper. All authors reviewed, revised and approved the manuscript for publication.

## Acknowledgements

The authors thank Jeroen Drenth for his helpful reading and suggestions on the manuscript. MLM and MJA are members of the Researcher Career from CONICET, Argentina.

## References

1. Cheeseman, P., A. Toms-Wood, and R.S. Wolfe, Isolation and Properties of a Fluorescent Compound, Factor 420, from Methanobacterium Strain M.o.H. Journal of Bacteriology, 1972. 112(1): p. 527–531.

2. Eirich, L.D., G.D. Vogels, and R.S. Wolfe, Proposed structure for coenzyme F420 from methanobacterium. Biochemistry, 1978. 17(22): p. 4583–4593.

3. Greening, C., et al., Physiology, Biochemistry, and Applications of F420- and Fo-Dependent Redox Reactions. Microbiol Mol Biol Rev, 2016. 80(2): p. 451–93.

4. Zehnder, A.J.B., et al., Characterization of an acetate-decarboxylating, non-hydrogen-oxidizing methane bacterium. Archives of Microbiology, 1980. 124(1): p. 1–11.

5. Belay, N., R. Sparling, and L. Daniels, Relationship of formate to growth and methanogenesis by Methanococcus thermolithotrophicus. Applied and Environmental Microbiology, 1986. 52(5): p. 1080–1085.

6. Jirapanjawat, T., et al., The redox cofactor F420 protects mycobacteria from diverse antimicrobial compounds and mediates a reductive detoxification system. Applied and environmental microbiology, 2016. 82(23): p. 6810–6818.

7. Selengut, J.D. and D.H. Haft, Unexpected abundance of coenzyme F420-dependent enzymes in Mycobacterium tuberculosis and other actinobacteria. Journal of bacteriology, 2010. 192(21): p. 5788–5798.

8. Campbell, Z.T., et al., Crystal structure of the bacterial luciferase/flavin complex provides insight into the function of the beta subunit. Biochemistry, 2009. 48(26): p. 6085–94.

9. Mashalidis, E.H., et al., Rv2607 from Mycobacterium tuberculosis Is a Pyridoxine 50-Phosphate Oxidase with Unusual Substrate Specificity. PLOS ONE, 2011. 6(11): p. e27643.

10. Lapalikar, G.V., et al., F420H2-Dependent Degradation of Aflatoxin and other Furanocoumarins Is Widespread throughout the Actinomycetales. PLOS ONE, 2012. 7(2): p. e30114.

11. Taylor, M.C., et al., Identification and characterization of two families of F420H2-dependent reductases from Mycobacteria that catalyse aflatoxin degradation. Molecular Microbiology, 2010. 78(3): p. 561–575.

12. Ahmed, F.H., et al., Sequence-Structure-Function Classification of a Catalytically Diverse Oxidoreductase Superfamily in Mycobacteria. J Mol Biol, 2015. 427(22): p. 3554–3571.

13. White, R.H., Biochemical Origins of Lactaldehyde and Hydroxyacetone in Methanocaldococcus jannaschii. Biochemistry, 2008. 47(17): p. 5037–5046.

14. Kumar, H., et al., Isolation and characterization of a thermostable F420:NADPH oxidoreductase from Thermobifida fusca. Journal of Biological Chemistry, 2017. 292(24): p. 10123–10130.

15. Seedorf, H., et al., F420H2 oxidase (FprA) from Methanobrevibacter arboriphilus, a coenzyme F420-dependent enzyme involved in O2 detoxification. Archives of Microbiology, 2004. 182(2): p. 126–137.

16. Hagemeier, C.H., et al., Coenzyme F420-dependent methylenetetrahydromethanopterin dehydrogenase (Mtd) from Methanopyrus kandleri: a methanogenic enzyme with an unusual quarternary structure. J Mol Biol, 2003. 332(5): p. 1047–57.

17. Vitt, S., et al., The F420-Reducing [NiFe]-Hydrogenase Complex from Methanothermobacter marburgensis, the First X-ray Structure of a Group 3 Family Member. Journal of Molecular Biology, 2014. 426(15): p. 2813–2826.

18. Susanti, D., U. Loganathan, and B. Mukhopadhyay, A Novel F420-dependent Thioredoxin Reductase Gated by Low Potential FAD: A TOOL FOR REDOX REGULATION IN AN ANAEROBE. Journal of Biological Chemistry, 2016. 291(44): p. 23084–23100.

19. NOMENCLATURE COMMITTEE OF THE INTERNATIONAL UNION OF BIOCHEMISTRY AND MOLECULAR BIOLOGY, in Enzyme Nomenclature. 1992, Academic Press: San Diego. p. ii.

20. Omelchenko, M.V., et al., Non-homologous isofunctional enzymes: a systematic analysis of alternative solutions in enzyme evolution. Biology direct, 2010. 5: p. 31–31.

21. Orengo, C.A. and J.M. Thornton, PROTEIN FAMILIES AND THEIR EVOLUTION—A STRUCTURAL PERSPECTIVE. Annual Review of Biochemistry, 2005. 74(1): p. 867–900.

22. Harms, M.J. and J.W. Thornton, Evolutionary biochemistry: revealing the historical and physical causes of protein properties. Nature Reviews Genetics, 2013. 14(8): p. 559–571.

23. Dawson, N.L., et al., CATH: an expanded resource to predict protein function through structure and sequence. Nucleic Acids Research, 2016. 45(D1): p. D289–D295.

24. Aufhammer, S.W., et al., Crystal structure of methylenetetrahydromethanopterin reductase (Mer) in complex with coenzyme F420: Architecture of the F420/FMN binding site of enzymes within the nonprolyl cis-peptide containing bacterial luciferase family. Protein Sci, 2005. 14(7): p. 1840–9.

25. Bashiri, G., et al., Crystal structures of F420-dependent glucose-6-phosphate dehydrogenase FGD1 involved in the activation of the anti-tuberculosis drug candidate PA-824 reveal the basis of coenzyme and substrate binding. J Biol Chem, 2008. 283(25): p. 17531–41.

26. Aufhammer, S.W., et al., Coenzyme binding in F420-dependent secondary alcohol dehydrogenase, a member of the bacterial luciferase family. Structure, 2004. 12(3): p. 361–70.

27. Eichhorn, E., et al., Crystal structure of Escherichia coli alkanesulfonate monooxygenase SsuD. J Mol Biol, 2002. 324(3): p. 457–68.

28. Fisher, A.J., et al., The 1.5-Å Resolution Crystal Structure of Bacterial Luciferase in Low Salt Conditions. Journal of Biological Chemistry, 1996. 271(36): p. 21956–21968.

29. Mascotti, M.L., et al., Reconstructing the evolutionary history of F420-dependent dehydrogenases. Scientific Reports, 2018. 8(1): p. 17571.

30. Warkentin, E., et al., Structures of F420-H2:NADP+ oxidoreductase with and without its substrates bound. The EMBO Journal, 2001. 20(23): p. 6561.

31. Seedorf, H., et al., Structure of coenzyme F420H2 oxidase (FprA), a di-iron flavoprotein from methanogenic Archaea catalyzing the reduction of O2 to H2O. The FEBS Journal, 2007. 274(6): p. 1588–1599.

32. Nocek, B., et al., Crystal Structures of Δ1-Pyrroline-5-carboxylate Reductase from Human Pathogens Neisseria meningitides and Streptococcus pyogenes. Journal of Molecular Biology, 2005. 354(1): p. 91–106.

33. Carter, M.S., et al., Functional assignment of multiple catabolic pathways for d-apiose. Nature Chemical Biology, 2018. 14(7): p. 696–705.

34. Mukhopadhyay, B., et al., Cloning, sequencing, and transcriptional analysis of the coenzyme F420-dependent methylene-5,6,7,8-tetrahydromethanopterin dehydrogenase gene from Methanobacterium thermoautotrophicum strain Marburg and functional expression in Escherichia coli. The Journal of biological chemistry, 1995. 270(6): p. 2827–2832.

35. Schwörer, B., et al., Formylmethanofuran: tetrahydromethanopterin formyltransferase and N5,N10-methylenetetrahydromethanopterin dehydrogenase from the sulfate-reducing Archaeoglobus fulgidus: similarities with the enzymes from methanogenic Archaea. Archives of microbiology, 1993. 159(3): p. 225–232.

36. Klenk, H.-P., et al., The complete genome sequence of the hyperthermophilic, sulphate-reducing archaeon Archaeoglobus fulgidus. Nature, 1997. 390(6658): p. 364–370.

37. Silaghi-Dumitrescu, R., et al., X-ray Crystal Structures of Moorella thermoacetica FprA. Novel Diiron Site Structure and Mechanistic Insights into a Scavenging Nitric Oxide Reductase. Biochemistry, 2005. 44(17): p. 6492–6501.

38. Hillmann, F., et al., Reductive dioxygen scavenging by flavo-diiron proteins of Clostridium acetobutylicum. FEBS letters, 2009. 583(1): p. 241–245.

39. Silaghi-Dumitrescu, R., et al., A Flavodiiron Protein and High Molecular Weight Rubredoxin from Moorella thermoacetica with Nitric Oxide Reductase Activity. Biochemistry, 2003. 42(10): p. 2806–2815.

40. Cellitti, Susan E., et al., Structure of Ddn, the Deazaflavin-Dependent Nitroreductase from Mycobacterium tuberculosis Involved in Bioreductive Activation of PA-824. Structure. 20(1): p. 101–112.

41. Kim, S.-H., et al., Crystal structure of the flavin reductase component (HpaC) of 4-hydroxyphenylacetate 3-monooxygenase from Thermus thermophilus HB8: Structural basis for the flavin affinity. Proteins: Structure, Function, and Bioinformatics, 2008. 70(3): p. 718–730.

42. van den Heuvel, R.H.H., et al., Structural Studies on Flavin Reductase PheA2 Reveal Binding of NAD in an Unusual Folded Conformation and Support Novel Mechanism of Action. Journal of Biological Chemistry, 2004. 279(13): p. 12860–12867.

43. Duffner, F.M., et al., Phenol/cresol degradation by the thermophilic Bacillus thermoglucosidasius A7: cloning and sequence analysis of five genes involved in the pathway. Gene, 2000. 256(1): p. 215–221.

44. Lapalikar, G.V., et al., Cofactor promiscuity among F420-dependent reductases enables them to catalyse both oxidation and reduction of the same substrate. Catalysis Science & Technology, 2012. 2(8): p. 1560–1567.

45. Mills, D.J., et al., De novo modeling of the F420-reducing [NiFe]-hydrogenase from a methanogenic archaeon by cryo-electron microscopy. eLife, 2013. 2: p. e00218.

46. Ilina, Y., et al., X-ray Crystallography and Vibrational Spectroscopy Reveal the Key Determinants of Biocatalytic Dihydrogen Cycling by [NiFe] Hydrogenases. Angewandte Chemie International Edition, 2019. 58(51): p. 18710–18714.

47. Welte, C. and U. Deppenmeier, Re-evaluation of the function of the F420 dehydrogenase in electron transport of Methanosarcina mazei. The FEBS Journal, 2011. 278(8): p. 1277–1287.

48. Johnson, E.F. and B. Mukhopadhyay, Coenzyme F_420_-Dependent Sulfite Reductase-Enabled Sulfite Detoxification and Use of Sulfite as a Sole Sulfur Source by *Methanococcus maripaludis*. Applied and Environmental Microbiology, 2008. 74(11): p. 3591–3595.

49. Yan, Z., M. Wang, and J.G. Ferry, A Ferredoxin- and F420H2-Dependent, Electron-Bifurcating, Heterodisulfide Reductase with Homologs in the Domains Bacteria and Archaea. mBio, 2017. 8(1): p. e02285–16.

50. Susanti, D., et al., Thioredoxin targets fundamental processes in a methane-producing archaeon, Methanocaldococcus jannaschii. Proceedings of the National Academy of Sciences, 2014. 111(7): p. 2608–2613.

51. Laskowski, R.A., et al., PDBsum: Structural summaries of PDB entries. Protein Science, 2018. 27(1): p. 129–134.

52. El-Gebali, S., et al., The Pfam protein families database in 2019. Nucleic Acids Research, 2018. 47(D1): p. D427–D432.

53. Mitchell, A.L., et al., InterPro in 2019: improving coverage, classification and access to protein sequence annotations. Nucleic Acids Research, 2018. 47(D1): p. D351–D360.

54. Pei, J., B.-H. Kim, and N.V. Grishin, PROMALS3D: a tool for multiple protein sequence and structure alignments. Nucleic Acids Research, 2008. 36(7): p. 2295–2300.

55. Katoh, K., J. Rozewicki, and K.D. Yamada, MAFFT online service: multiple sequence alignment, interactive sequence choice and visualization. Briefings in Bioinformatics, 2017. 20(4): p. 1160–1166.

56. Fu, L., et al., CD-HIT: accelerated for clustering the next-generation sequencing data. Bioinformatics, 2012. 28(23): p. 3150–3152.

57. Darriba, D., et al., ProtTest 3: fast selection of best-fit models of protein evolution. Bioinformatics, 2011. 27(8): p. 1164–1165.

58. Guindon, S., et al., New Algorithms and Methods to Estimate Maximum-Likelihood Phylogenies: Assessing the Performance of PhyML 3.0. Systematic Biology, 2010. 59(3): p. 307–321.

59. Stamatakis, A., RAxML version 8: a tool for phylogenetic analysis and post-analysis of large phylogenies. Bioinformatics, 2014. 30(9): p. 1312–1313.

60. Lemoine, F., et al., Renewing Felsenstein’s phylogenetic bootstrap in the era of big data. Nature, 2018. 556(7702): p. 452–456.

61. Mashalidis, E.H., et al., Molecular insights into the binding of coenzyme F420 to the conserved protein Rv1155 from Mycobacterium tuberculosis. Protein Science, 2015. 24(5): p. 729–740.

