## Supplementary material for "On the diversity of F_420_-dependent oxidoreductases: a sequence- and structure-based classification": Table S1

| Superfamily | PDB | Name | Type | Uniprot/Genbank |
|---|---|---|---|---|
| I | **1ezw** | MER_METKA | MER | Q8TXY4 |
| | **1f07** | MER_METTM | MER | Q50744 |
| | **1z69** | MER_METBF | MER | Q46FV4 |
| | **3b4y, 3c8n** | FGD_MYCTU | FGD | P9WNE1 |
| | **5lxe** | FGD1_RHOJR | FGD | Q0RVH7 |
| | **1rhc** | ADF | ADH | O93734 |
| II | **1jax, 1jay** | FNO_ARCFU | FNO | O29370 |
| | **2raf** | oxidoreductase | FNO | F9URQ8 |
| | **3dtt** | NADP oxidoreductase | FNO | A0JSJ2 |
| | **5n2i** | $F_{420}$:NADP oxidoreductase | FNO | Q47RA9 |
| | **1qv9,1u6i, 1u6k, 3iqe, 3iqf, 3iqz** | MTD_METKA | MTD | P94951 |
| | **2ohh, 2ohi, 2ohj** | FPRA_METTM | FPRA | Q50497 |
| | **6frm, 6frn** | $F_{420}H_2$ oxidase (FprA) | FRPA | WP_018154036.1 |
| III | **3f7e** | Msmeg_3380 | FDOR-B | A0QXP8 |
| | **3h96** | Msmeg_3358 $F_{420}$ reductase | FDOR-A | A0QXM5 |
| | **3r5r, 3r5w** | DDN_MYCTU | DDN | P9WP15 |
| | **3r5y** | [redacted] | [redacted] | Q5YUF0 |
| | **3r5z** | [redacted] | [redacted] | Q5YYT7 |
| | **4qvb** | Rv1155 | FDOR-B | O06553 |
| | **4y9i** | $F_{420}H_2$ dependent reductase (fdr-a) | FDOR-A | A0QU01 |
| | **4zky, 5jv4** | Msmeg_6526 | FDOR-B | A0R6F1 |
| | **5jab** | biliverdin reductase Rv2074 | FDOR-B | P9WLL7 |
| IV | **4omf, 3zfs, 4ci0** | $F_{420}$-reducing [NiFe]-hydrogenase, subunit beta | FrhABG | D9PYF6 |
| | **6qgr, 6qgt, 6gii** | $F_{420}$-reducing [NiFe]-hydrogenase, subunit beta | FrhABG | A0A0E3QWH3 |
| | [redacted] | | Fpo-F | Q8PZ67 |

| | | | | |
|---|---|---|---|---|
| | [redacted] | Coenzyme $F_{420}$-dependent sulfite reductase | Fsr (N-term) | Q58280 |
| **V** | [redacted] | $F_{420}$-dependent thioredoxin reductase | DFTR | Q58931 |

| Organism | Lenght | PFAM | | CATH | |
|---|---|---|---|---|---|
| *Methanopyrus kandleri* | 347 | PF00296 | Bac_luciferase | 3.20.20.30 | Alpha-Beta Barrel |
| *Methanobacterium thermoautotrophicum* | 321 | | | | |
| *Methanosarcina barkeri* | 327 | | | | |
| *mycobacterium tuberculosis* | 332 | | | | |
| *Rhodococcus jostii* | 324 | | | | |
| *Methanoculleus thermophilus* | 330 | | | | |
| *Archaeoglobus fulgidus* | 212 | PF03807 | F420_oxidored | 3.40.50.720 | 3-Layer(aba) Sandwich |
| *Lactobacillus plantarum* | 190 | | | | |
| *Arthrobacter sp* | 220 | | | | |
| *Thermobifida fusca* | 225 | | | | |
| *Methanopyrus kandleri* | 282 | PF01993 | MTD | 3.40.50.10830 | 3-Layer(aba) Sandwich |
| *Methanothermobacter marburgensis* | 403 | PF00753 | Lactamase_B | 3.60.15.10 | 4-Layer Sandwich |
| | | PF00258 | Flavodoxin_1 | 3.40.50.360 | 3-Layer(aba) Sandwich |
| *Methanothermococcus thermolithotrophicus* | 402 | PF00753 | Lactamase_B | 3.60.15.10 | 4-Layer Sandwich |
| | | PF00258 | Flavodoxin_1 | 3.40.50.360 | 3-Layer(aba) Sandwich |
| *Mycobacterium smegmatis* | 128 | PF01243 | Putative_PNPOx | 2.30.110.10 | Mainly Beta Roll |
| *Mycobacterium smegmatis* | 137 | | | | |
| *Mycobacterium tuberculosis* | 109 | | | | |
| *Nocardia farcinica* | 141 | | | | |
| *Nocardia farcinica* | 140 | | | | |
| *Mycobacterium tuberculosis* | 141 | | | | |
| *Mycobacterium smegmatis* | 114 | PF04075 | F420H2_quin_red | | |
| *Mycobacterium smegmatis* | 139 | PF01243 | Putative_PNPOx | | |
| *Mycobacterium tuberculosis* | 135 | | | | |
| *Methanothermobacter marburgensis* | 281 | PF04432 PF04422 | Coenzyme $F_{420}$ hydrogenase/ dehydrogenase, beta [illegible] | 2.30.30.1030 | SH3 barrel |
| *Methanosarcina barkeri* | 291 | | | | |
| *Methanosarcina mazei* | 346 | | | | |

| | | | | | |
|---|---|---|---|---|---|
| *Methanocaldococcus jannaschii* | 610 (310 N-term) | | subunit N-term/C-term | | |
| *Methanocaldococcus jannaschii* | 301 | PF07992 | Pyr_redox_2 | 3.50.50.60 | FAD/NAD(P)-binding domain |

| Reference |
| --- |
| Shima et al, J Mol Biol (2000) 300(4):935-950 |
| Aufhammer et al, Prot Sci (2005) 14(7):1840-1849 |
| Bashiri et al, J Biol Chem (2008) 283:17531-17541 |
| Nguyen et al, Appl Microbiol Biotechnol (2017) 101:2831-2842 |
| Aufhammer et al, Structure (2004) 12:361-370 |
| Warkentin et al, EMBO J (2001) 20(23):6561 |
| [redacted] |
| Kumar et al, J Biol Chem (2017) 292(24):10123-10130 |
| Hagemeier et al, J Mol Biol (2003) 332:1047-1057 |
| Seedorf et al, FEBS J (2007) **274**(6):1588-1599 |
| Engilberge et al, Chemistry (2018) 24(39):9739-9746 |
| [redacted] |
| Taylor et al, Mol Microbiol (2010) 78:561-575 |
| Celliti et al, Structure (2012) 20:101-112 |
| Mashalidis et al, Protein Sci (2015)24:729-740 |
| Ahmed et al, J Mol Biol (2015)427:3554-3571 |
| Ahmed et al, Protein Sci (2016)25:1692-1709 |
| Vit et al, J Mol Biol (2014) 426:2813-2826 |
| Mills et al, eLife (2013) 2:e00218 |
| Ilina et al, Angew Chem Int Ed (2019) 58:18710-18714 |
| Welte & Deppenmeier, FEBS J (2011) 278:1277-1287 |

| |
|---|
| Johnson & Mukhopadhyay, J Biol Chem (2005) 280(46):38776-38786 |
| Susanti et al, J Biol Chem (2016) 291(44):23084-23100 |
