## Supplementary material for "On the diversity of F_420_-dependent oxidoreductases: a sequence- and structure-based classification": Fig. S1, Fig. S2, Fig. S3, Fig. S4

### Supporting information

---

**Figure S1**

### *Superfamily I: TIM barrel $F_{420}$-dependent enzymes*

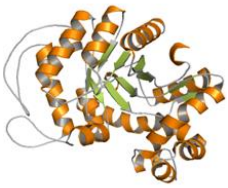

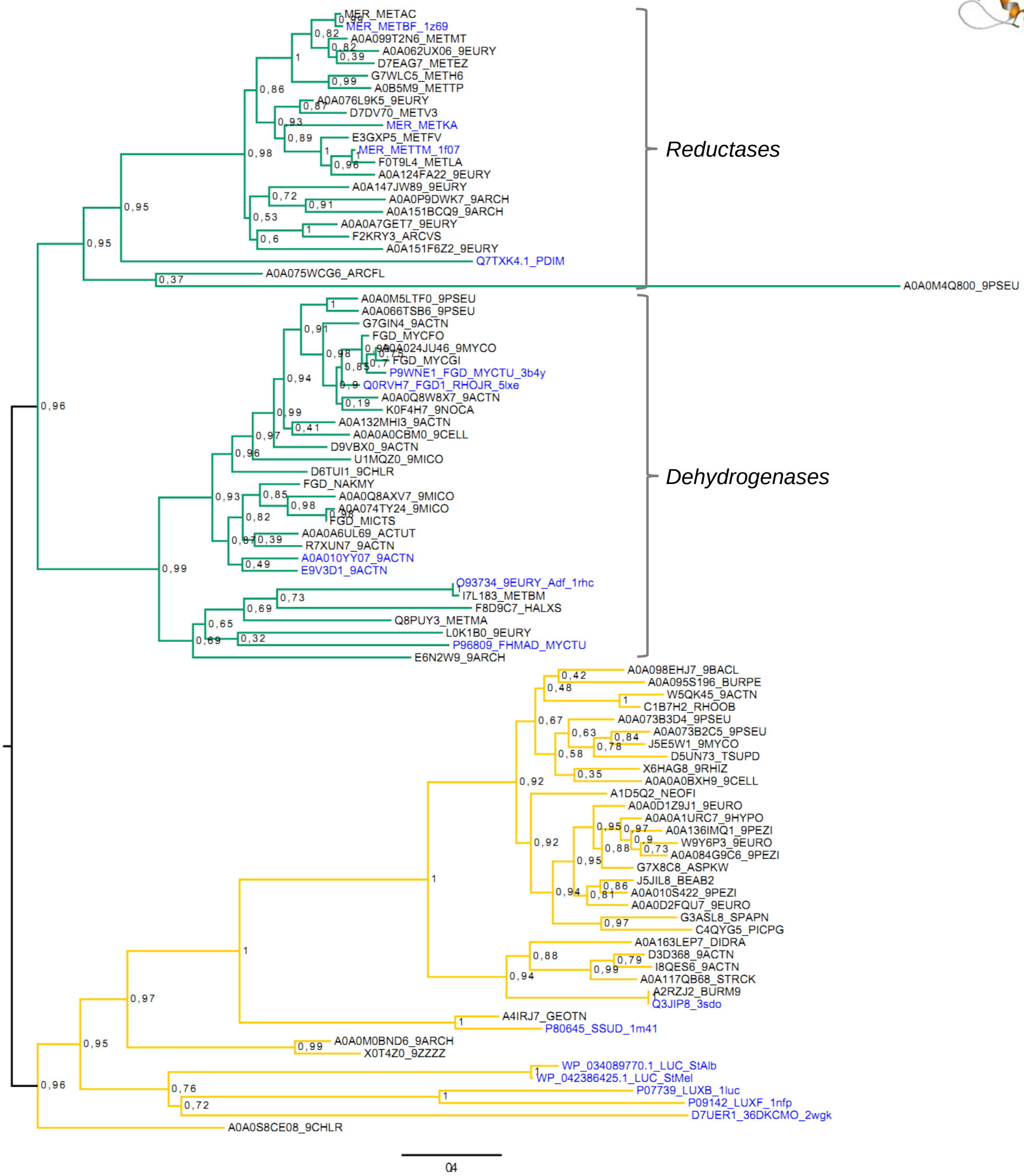


**Phylogeny of Superfamily I. Sequence-based tree.** The tree was constructed in RaxML, 500 BS were performed and subjected to bootstrap transfer. TBE values are shown at the nodes. Rooting was performed according to Mascotti *et al*[1]. The MSA contained 91 sequences (max. identity 80%) and 311 sites. Uniprot ID_name (when available)_taxonomy_PDB ID (when available) is given for each sequence. All enzymes experimentally characterized are shown in blue. Colored branches indicate the cofactor specificity: $F_{420}$-dependent enzymes (green) and FMN-using enzymes (yellow).

[1] M.L. Mascotti, H. Kumar, Q-T. Nguyen, M.J. Ayub, M.W. Fraaije, Reconstructing the evolutionary history of $F_{420}$-dependent dehydrogenases, Scientific Reports 8 (2018) 17571. 10.1038/s41598-018-35590-2.

**Figure S2**

### *Superfamily II: Rossmann fold $F_{420}$-dependent enzymes*

**A)**

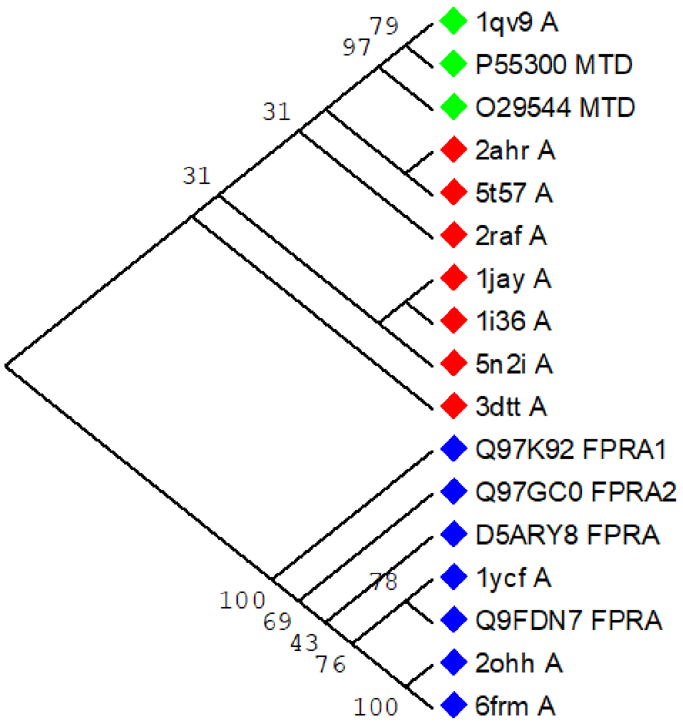


**Phylogenies of families belonging to Superfamily II**
**A) Structure-based tree.** The structural alignment was obtained in PROMALS3D (17 structures, 240 sites). The tree was constructed in PhyML, 100 BS were run and values > 50 are shown at the nodes. Tree is shown as midpoint rooted. Families are marked with colored diamonds as follows: FNOs (red), MTDs (green) and oxidases (blue).

B)

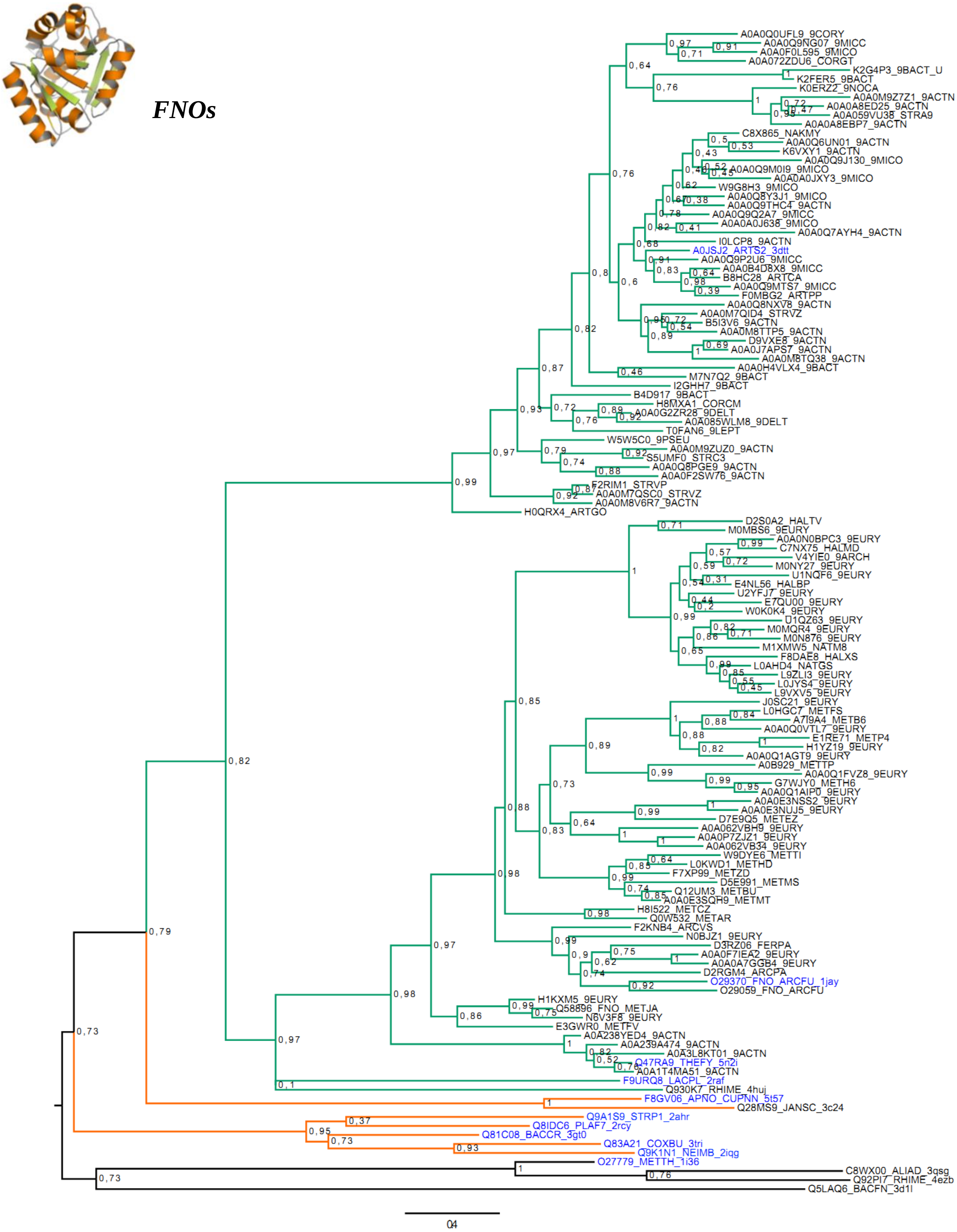

***FNOs***

A0A0Q0UFL9_9CORY
A0A0Q9NG07_9MICC
A0A0F0L595_9MICO
A0A072ZDU6_CORGT
K2G4P3_9BACT_U
K2FER5_9BACT
K0ERZ2_9NOCA
A0A0M9Z7Z1_9ACTN
A0A0A8ED25_9ACTN
A0A059VU38_STRA9
A0A0A8EBP7_9ACTN
C8X865_NAKMY
A0A0Q6UN01_9ACTN
K6VXY1_9ACTN
A0A0Q9J130_9MICO
A0A0Q9M0I9_9MICO
A0A0A0JXY3_9MICO
W9G8H3_9MICO
A0A0Q8Y3J1_9MICO
A0A0Q9THC4_9ACTN
A0A0Q9Q2A7_9MICC
A0A0A0J638_9MICO
A0A0Q7AYH4_9ACTN
I0LCP8_9ACTN
A0JSJ2_ARTS2_3dtt
A0A0Q9P2U6_9MICC
A0A0B4D8X8_9MICC
B8HC28_ARTCA
A0A0Q9MTS7_9MICC
F0MBG2_ARTPP
A0A0Q8NXV8_9ACTN
A0A0M7QID4_STRVZ
B5I3V6_9ACTN
A0A0M8TTP5_9ACTN
D9VXE8_9ACTN
A0A0J7APS7_9ACTN
A0A0M8TQ38_9ACTN
A0A0H4VLX4_9BACT
M7N7Q2_9BACT
I2GHH7_9BACT
B4D917_9BACT
H8MXA1_CORCM
A0A0G2ZR28_9DELT
A0A085WLM8_9DELT
T0FAN6_9LEPT
W5W5C0_9PSEU
A0A0M9ZUZ0_9ACTN
S5UMF0_STRC3
A0A0Q8PGE9_9ACTN
A0A0F2SW76_9ACTN
F2RIM1_STRVP
A0A0M7QSC0_STRVZ
A0A0M8V6R7_9ACTN
H0QRX4_ARTGO
D2S0A2_HALTV
M0MBS6_9EURY
A0A0N0BPC3_9EURY
C7NX75_HALMD
V4YIE0_9ARCH
M0NY27_9EURY
U1NQF6_9EURY
E4NL56_HALBP
U2YFJ7_9EURY
E7QU00_9EURY
W0K0K4_9EURY
U1QZ63_9EURY
M0MQR4_9EURY
M0N876_9EURY
M1XMW5_NATM8
F8DAE8_HALXS
L0AHD4_NATGS
L9ZLI3_9EURY
L0JYS4_9EURY
L9VXV5_9EURY
J0SC21_9EURY
L0HGC7_METFS
A7I9A4_METB6
A0A0Q0VTL7_9EURY
E1RE71_METP4
H1YZ19_9EURY
A0A0Q1AGT9_9EURY
A0B929_METTP
A0A0Q1FVZ8_9EURY
G7WJY0_METH6
A0A0Q1AIP0_9EURY
A0A0E3NSS2_9EURY
A0A0E3NUJ5_9EURY
D7E9Q5_METEZ
A0A062VBH9_9EURY
A0A0P7ZJZ1_9EURY
A0A062VB34_9EURY
W9DYE6_METTI
L0KWD1_METHD
F7XP99_METZD
D5E991_METMS
Q12UM3_METBU
A0A0E3SQH9_METMT
H8I522_METCZ
Q0W532_METAR
F2KNB4_ARCVS
N0BJZ1_9EURY
D3RZ06_FERPA
A0A0F7IEA2_9EURY
A0A0A7GGB4_9EURY
D2RGM4_ARCPA
O29370_FNO_ARCFU_1jay
O29059_FNO_ARCFU
H1KXM5_9EURY
Q58896_FNO_METJA
N6V3F8_9EURY
E3GWR0_METFV
A0A238YED4_9ACTN
A0A239A474_9ACTN
A0A3L8KT01_9ACTN
Q47RA9_THEFY_5n2i
A0A1T4MA51_9ACTN
F9URQ8_LACPL_2raf
Q930K7_RHIME_4huj
F8GV06_APNO_CUPNN_5t57
Q28MS9_JANSC_3c24
Q9A1S9_STRP1_2ahr
Q8IDC6_PLAF7_2rcy
Q81C08_BACCR_3gt0
Q83A21_COXBU_3tri
Q9K1N1_NEIMB_2iqg
O27779_METTH_1i36
C8WX00_ALIAD_3qsg
Q92PI7_RHIME_4ezb
Q5LAQ6_BACFN_3d1l
04

**C)**

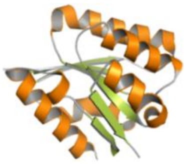

***MTDs***

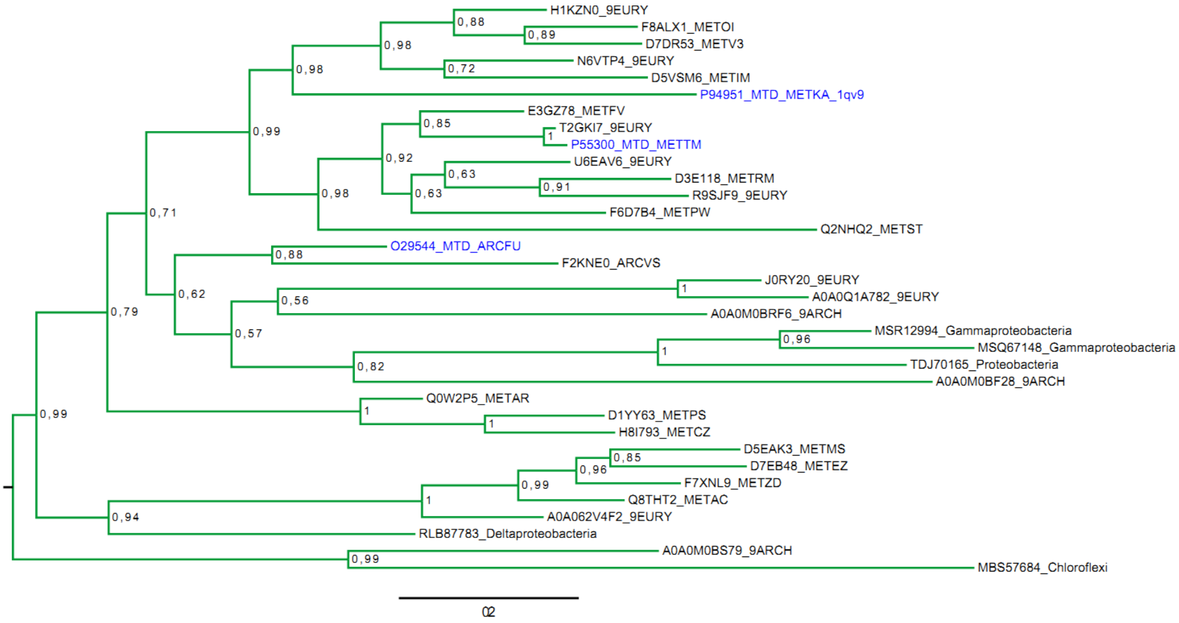
H1KZN0_9EURY
F8ALX1_METOI
D7DR53_METV3
N6VTP4_9EURY
D5VSM6_METIM
P94951_MTD_METKA_1qv9
E3GZ78_METFV
T2GKI7_9EURY
P55300_MTD_METTM
U6EAV6_9EURY
D3E118_METRM
R9SJF9_9EURY
F6D7B4_METPW
Q2NHQ2_METST
O29544_MTD_ARCFU
F2KNE0_ARCVS
J0RY20_9EURY
A0A0Q1A782_9EURY
A0A0M0BRF6_9ARCH
MSR12994_Gammaproteobacteria
MSQ67148_Gammaproteobacteria
TDJ70165_Proteobacteria
A0A0M0BF28_9ARCH
Q0W2P5_METAR
D1YY63_METPS
H8I793_METCZ
D5EAK3_METMS
D7EB48_METEZ
F7XNL9_METZD
Q8THT2_METAC
A0A062V4F2_9EURY
RLB87783_Deltaproteobacteria
A0A0M0BS79_9ARCH
MBS57684_Chloroflexi
0,88
0,89
0,98
0,72
0,98
0,99
0,85
1
0,92
0,63
0,91
0,63
0,98
0,71
0,88
0,62
1
0,56
0,57
0,96
1
0,82
0,79
1
1
0,99
0,85
0,96
0,99
1
0,94
0,99

D)

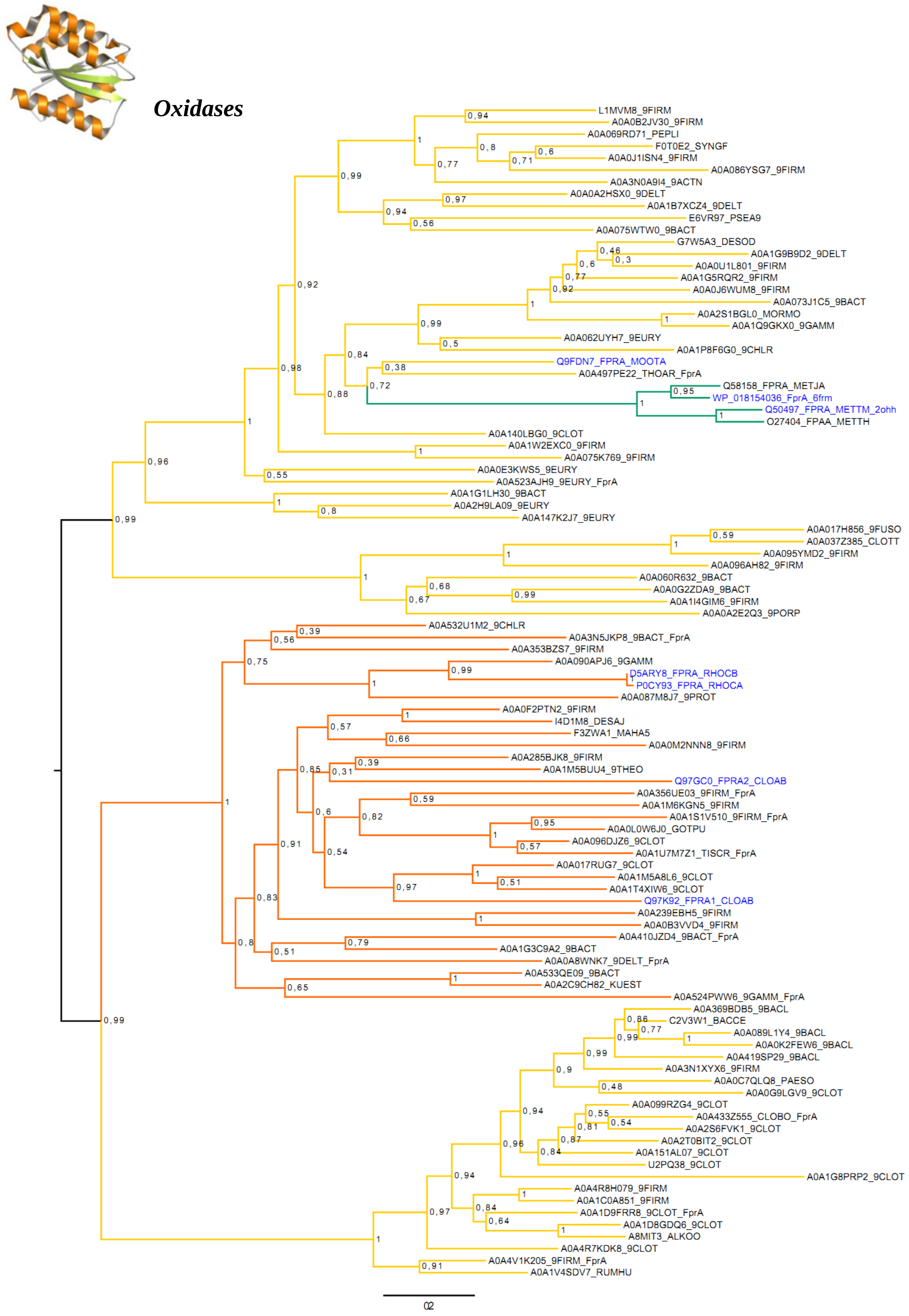


**B,C,D) Sequence-based trees.** The trees were constructed in RaxML, 500 BS were performed and subjected to bootstrap transfer. TBE values are shown at the nodes. Midpoint rooting was used in all cases. Uniprot ID_name (when available)_taxonomy_PDB ID (when available) is given for each sequence. All enzymes experimentally characterized are shown in blue. Colored branches indicate the cofactor specificity:

$F_{420}$-dependent enzymes (green), FMN-using enzymes (yellow) and NADPH using enzymes (orange). Unknown cofactor preference is indicated with black branches. **B)** FNOs family. The MSA contained 129 sequences (max. identity 80%) and 222 sites. **C)** MTDs family. The MSA contained 33 sequences (max id 80%) and 275 sites. **D)** Oxidases family. The MSA contained 91 sequences (max id 80%) and 391 sites. All enzymes in this family are inferred to be FMN-dependent and show different preference for the electron donor.

**Figure S3**

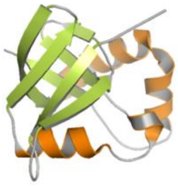

### *Superfamily III: β-roll $F_{420}$-dependent enzymes*

**A)**

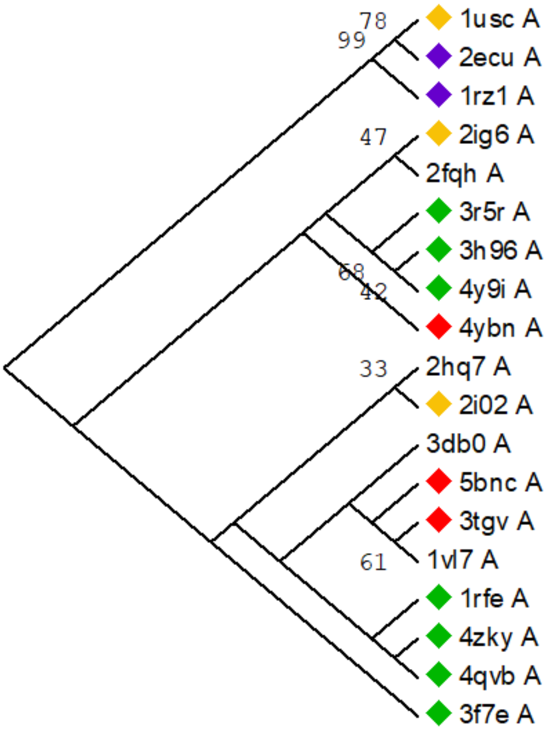


**Phylogeny of Superfamily III**

**A) Structure-based tree.** The structural alignment was obtained in PROMALS3D (19 structures, 128 sites). The tree was constructed in PhyML, 100BS were run and values > 50 are shown at the nodes. Tree is shown as midpoint rooted. Cofactor preference is indicated with colored diamonds as follows: $F_{420}$ (green), FMN (yellow), FAD (violet) and heme (red).

**B)**

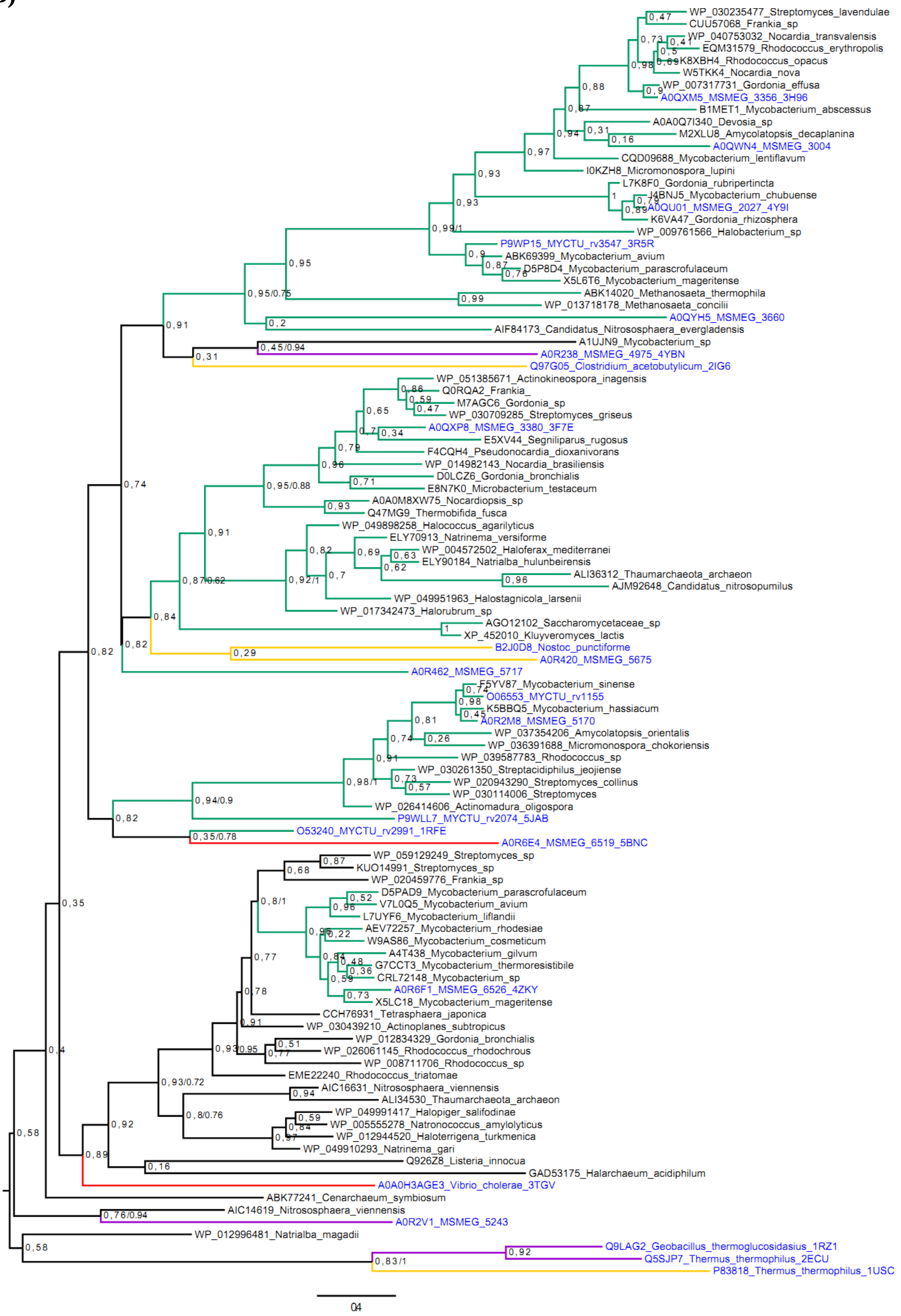


**B) Sequence-based tree.** The tree was constructed in RaxML, 500 BS were performed and subjected to bootstrap transfer. TBE values are shown at the nodes. Midpoint rooting was used. The MSA contained 104 sequences (max. identity 80%) and 156 sites. Phylogeny was also inferred in Mr. Bayes and the posterior probability values (PP) corresponding to major clade divergences in the tree are shown next to the TBE values. Uniprot ID_name (when available)_taxonomy_PDB ID (when available) is given for each sequence. All enzymes experimentally characterized are shown in blue. Colored branches indicate the cofactor specificity: $F_{420}$-dependent enzymes (green), FMN-using enzymes (yellow), FAD-using enzymes (violet) and heme-binding enzyme (red). Unknown cofactor preference is indicated with black branches.

**Figure S4**

### *Superfamily IV: SH3 barrel $F_{420}$-dependent enzymes*

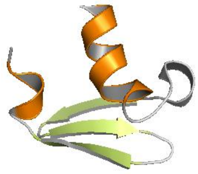

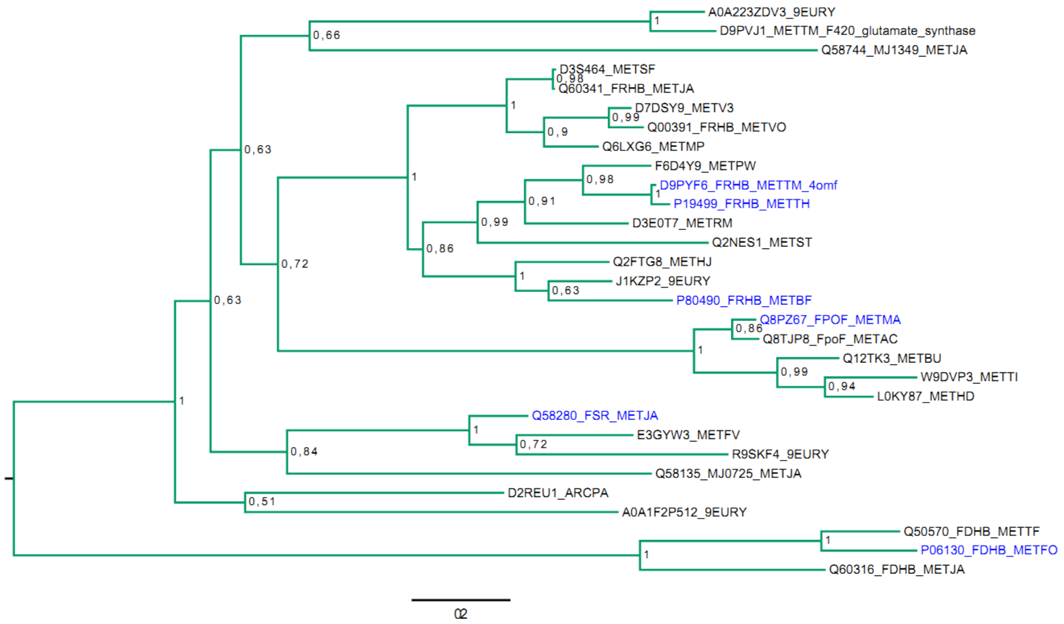


**Phylogeny of Superfamily IV. Sequence-based tree**. The tree was constructed in RaxML, 500 BS were performed and subjected to bootstrap transfer. TBE values are shown at the nodes. Midpoint rooting was used. The MSA contained 30 sequences (max. identity 80%) and 274 sites. Uniprot ID_name (when available)_taxonomy_PDB ID (when available) is given for each sequence. All enzymes experimentally characterized are shown in blue. Green branches indicate the $F_{420}$ cofactor specificity.
